# EnTrance: Exploration of Entropy Scaling Ball Cover Search in Protein Sequences

**DOI:** 10.1101/2021.05.31.446458

**Authors:** Yoonjin Kim, Zhen Guo, Jeffrey A. Robertson, Benjamin Reidys, Ziyan Zhang, Lenwood S. Heath

## Abstract

Biological sequence alignment using computational power has received increasing attention as technology develops. It is important to predict if a novel DNA sequence is potentially dangerous by determining its taxonomic identity and functional characteristics through sequence identification. This task can be facilitated by the rapidly increasing amounts of biological data in DNA and protein databases thanks to the corresponding increase in computational and storage costs. Unfortunately, the growth in biological databases has caused difficulty in exploiting this information. EnTrance presents an approach that can expedite the analysis of this large database by employing entropy scaling. This allows scaling with the amount of entropy in the database instead of scaling with the absolute size of the database. Since DNA and protein sequences are biologically meaningful, the space of biological sequences demonstrates the structure exploited by entropy scaling. As biological sequence databases grow, taking advantage of this structure can be extremely beneficial for reducing query times. EnTrance, the entropy scaling search algorithm introduced here, accelerates the biological sequence search exemplified by tools such as BLAST. EnTrance does this by utilizing a two step search approach. In this fashion, EnTrance quickly reduces the number of potential matches before more exhaustively searching the remaining sequences. Tests of EnTrance show that this approach can lead to improved query times. However, constructing the required entropy scaling indices beforehand can be challenging. To improve performance, EnTrance investigates several ideas for accelerating index build time that supports entropy scaling searches. In particular, EnTrance makes full use of the concurrency features of Go language greatly reducing the index build time. Our results identify key tradeoffs and demonstrate that there is potential in using these techniques for sequence similarity searches. Finally, EnTrance returns more matches and higher percentage identity matches when compared with existing tools.

## Introduction

As the size of biological databases grows exponentially(1,2), the importance of DNA sequencing technology emerges. Precise and fast DNA sequencing technology is essential for biological threat classifications with immense dataset(3, 4). Traditional sequence similarity search tools such as BLAST (5) and DIAMOND (6) have been used to identify potential threats with linear complexity scales, but they suffer a heavy computational burden due to large size of the dataset. In EnTrance, we propose a fast and efficient sequence similarity search tool using a pre-processed length-based binning and entropy scaling algorithm for extensive biological sequence databases. There are several sequence clusterization methods of UniRef(7) dataset for the Non-Redundant protein database which clusters and indexes similar sequences at varying levels of sequence identity such as CD-HIT(8) and LinClust(9). A compression algorithm CaBLASTP (10) introduced a local similarity-based compression method in NCBI NR database. This compression is based on the common protein sequences and links from the common sequences to the location in the original database. Another entropy scaling algorithm (11) compressed Protein Data Bank (PDB) to a hierarchical clustering structure by means of high-dimensional protein structure feature vectors. Mash tool (12) calculated pairwise Mash distance between genomes efficiently by *k*-mer “bottom-*h*” MinHash sketches and then clustered all genomes in NCBI RefSeq based on Mash distance. EnTrance adapts length-based binning clusters to achieve evenly distributed bins to search input using parallelization and ball construction of sequences within length similarities. Entropy scaling algorithms were proposed and exploited to facilitate bioinformatics research (11, 13–15). The entropy scaling searching algorithm improves searching performances (14) by effectively sequence a reduced number of the potential bins in the database such as NCBI NR and Uniref. The goal of EnTrance(Figure 2) is to leverage the internal structure of the large datasets in the sequence similarity search domain and to reveal the underlying threat level by taxonomy and functional analysis. The structure of EnTrance provides its search database *indexes* for large amounts of publicly available biological databases with sets of balls curated with a representative sequence referred to as *ball center* where the data is stored as the index. A two-step search inspired by the entropy-scaling algorithm, MICA (13), is designed as searching all the ball centers in the *coarse query* to find a small set of candidate balls. Then executes *fine query* as an exhaustive search (wrapping BLAST or DIAMOND) in the candidate balls to find top hits. Finally, using the top hits and their metadata, taxonomy and potential biological threat for a query sequence can be determined. As shown in the second box of system diagram(Figure 2), EnTrance also utilizes length based bins in parallel to boost query efficiency and produce refined results.

**Fig. 1.**
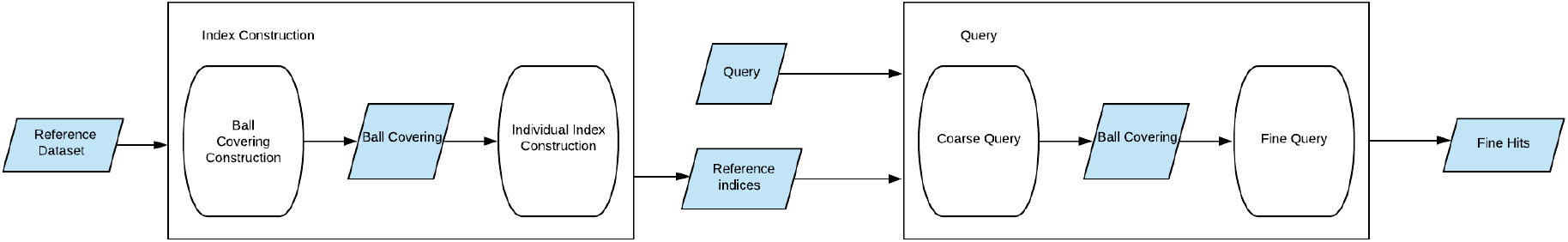
EnTrance system diagram without length based binning. It illustrates the system diagram of original design EnTrance where ball construction and query search were done in one database. EnTrance uses random ball construction to choose a fixed number of ball centers and generate ball covers on the dataset. When query search is requested, EnTrance calculates a distance between the input query and ball centers to find the nearest balls to continue fine search. Once the fine search is done, EnTrance sorts fine queries to output fine hits.

**Fig. 2.**
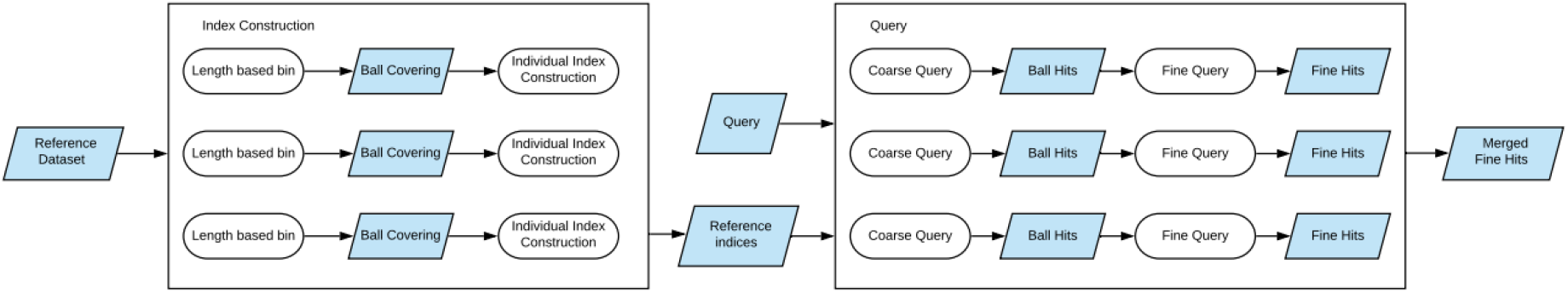
EnTrance system diagram with parallel length based binning search. Ball construction and coarse query search on each length based bins are done in parallel. Compared to the original diagram, new system pre-processes bins with length based binning and creates individual ball constructions with their own ball centers. Each length based bin has its own ball center from random ball construction and fine hits from fine search. EnTrance then merges fine hits from all length based bins.

EnTrance is primarily produced by Go language to supports lightweight concurrency and parallel system, which can efficiently speed up the index building process.

## System and Methods

### Entropy Scaling System

The premise of an entropy scaling algorithm is that some properties of the algorithm, such as execution time or storage cost, scale with the metric entropy of a dataset instead of with its size (13). The entropy scaling system in a large protein database *R* with a set of points *P* defines a *ball cover* as a set of balls *B* = {*B*_1_,*B*_2_,…,*B_n_*} where 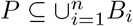 (Figure 3). Given a distance function *d* in a metric space, a *ball* is defined by a center point *c* and a radius *r*. The ball contains all points *p* in the space such that *d*(*c,p*) ≤ *r*. Entropy scaling approaches rely on the distance measure obeying the triangle inequality. If a similarity measure does not always obey the triangle inequality, these algorithms’ expected performance is proportional to the number of triplets in the data set for which the triangle inequality does hold (Supplemental Experimental Procedures and Figures from (13)). If *x* percent of the triplets in the set *T* = {{*a,b,c*} ⊆ *R*} obeys the triangle inequality, the similarity guarantees will be true approximately *x* percent of the time and the sensitivity of the search may also be affected.

**Fig. 3.**
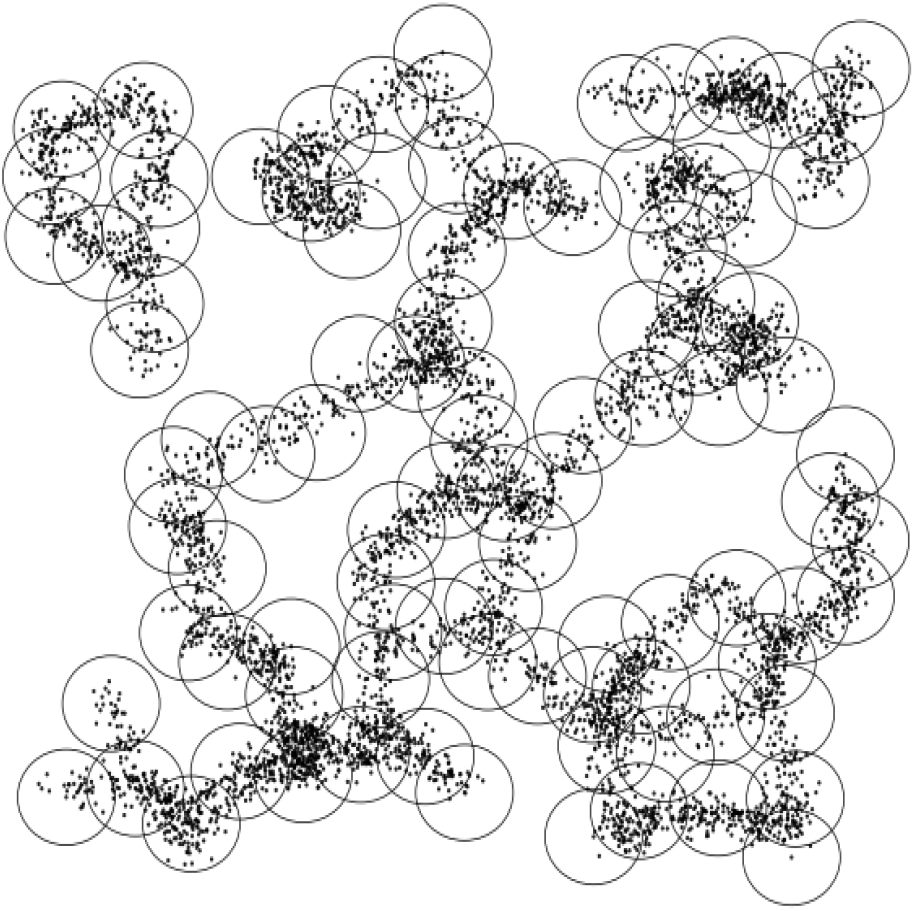
A ball covering of points in a 2D space

### Length based Binning

To create meaningful length-based bins we needed to ensure the bins have a dynamic range. This occurs because the sequence length distribution is heavily right skewed and most sequences have a length between 1-400 (Figure 4) and we wanted bins with a maximum of 1 million sequences. We decided on 1 million as the maximum number of sequences because the bins were wide enough for the 1-400 range and created enough bins outside the 1-400 range. This is important to keep in mind since we use these bins to generate ball covers for the later searches. The algorithm used to create the bins utilizes a greedy approach to bin creation. We create the bins sequentially by adding in all the sequences of a length until the bin reaches the maximum size. Once that occurs a new bin is created and we continue adding in sequences until all the sequences have been added.

#### Algorithm 1

*Sequence2Set* takes a sequence and an integer *k* and returns a set of all of the *k*-mers contained in the sequence

~~~
1: **procedure** Sequence2set(*Sequence S, int k*)
2:     *S* = [*s*_1_*s*_2_ ··· *S_n_*]
3:     *kmers* ← {}
4:     **for** *i* ← 1 **to** *n* – *k* **do**
5:         *kmers* = *kmers* ∪ {*s_i_* ··· *s*_*i*+*k*_}
6:     **return** *kmers*
~~~

**Fig. 4.**
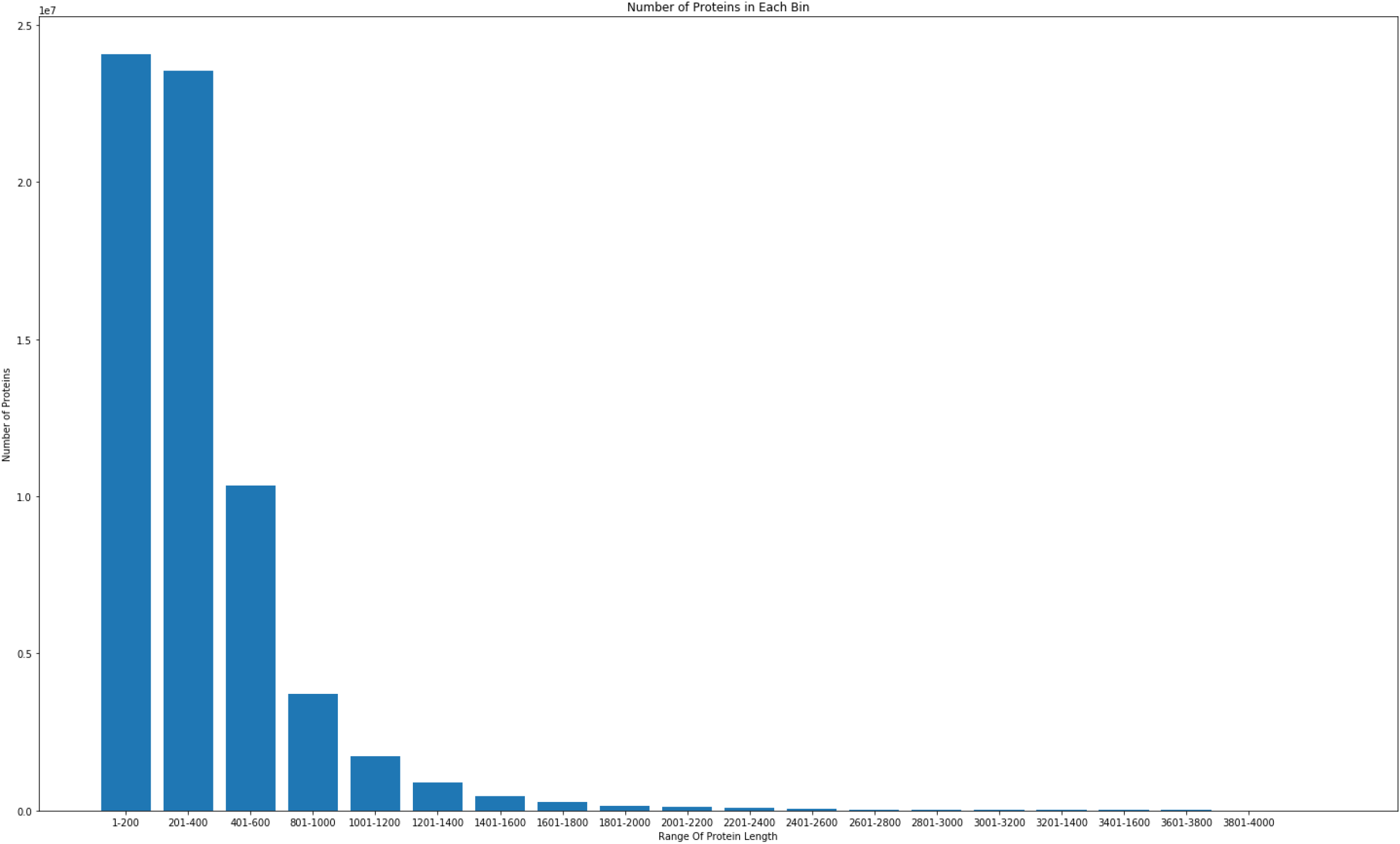
Distribution of protein lengths

### Jaccard Distance

Protein and DNA sequences can be regarded as a set of *k*-mers (Algorithm 1) which is a subsequence with k number of amino acids or nucleotides. With *k* = 2, for example, the sequence *S* = *AGTGGTC* is transformed into the set {*AG,GT,TG,GG,TC*} which is the set of all the 2-mers in *S*. Given two such sets, the Jaccard index (16) and Jaccard distance between them is defined as the number of elements that they have in common divided by the number of elements that are in either of them. A reduced alphabet where the number of protein alphabets is adapted into protein search to improve speeds and efficiency. In EnTrance, the protein alphabets are reduced to four letters using Murphy4 which maps 20 amino acids to 4({*L*: *LVIMC,A*: *AGSTP, F*: *FYW, E*: *EDNQKRH*}) following previous research (13). Formally, the Jaccard distance between sets *A* and *B* is

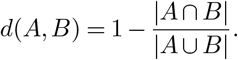

The value of the Jaccard distance varies from 0 (when the sets are identical) to 1 (when the sets disjoint). Jaccard distance can also be considered between multi-sets (17).

### MinHash Signature

MinHash (12, 18, 19) was specifically developed to speed up the approximation of Jaccard distance (16), although it can be heuristically used for other metrics. There are two types of MinHash signatures: bottom *h* values and *h* hashing functions. The bottom *h* method chooses a hashing function *H* and ranks the hashing values *H*(*S*) for all the *k*-mers *S* in a sequence. Then MinHash signature for *S* is the *h* least values in *H*(*S*). In *h* hashing functions method, a hashing function *H_i_*(0 < *i* ≤ *h*) can also be applied to each *k*-mer in *S* as *H_i_*(*S*). MinHash signature is composed of the minimum hashing value in *S* over each of the *h* hashing functions. Besides, the hashing functions and orders in *h* must be identical for all the sequences. First, a set of *h* permutations is created over the *k*-mer space. Each permutation is a deterministic ordering of all possible *k*-mers. Then, given a set of *k*-mers *S*, a MinHash signature (length *h*) of *S* is constructed by saving the location of minimum *k*-mer from *S* under each of the *h* permutations. The Hamming distance between two MinHash signatures approximates the Jaccard distance between the original *k*-mer sets.

### Evaluation Methods

These methods can be evaluated in several ways. Possibly the most important measures are query time, precision, and sensitivity (recall) (20). Precision is the fraction of returned sequences that are similar to the query, that is, the number of true positives divided by the total number of results. Sensitivity is the fraction of similar sequences that were in the index that were returned, that is, the number of true positives divided by the total number of sequences in the index which are similar to the query. Two other measures are index build time and index size. Because query time, index build time, and index size are all strongly related to the data set size, and this relationship could be dictated by the relative entropy of the data set, it is important that all methods be tested on data sets of significantly differing sizes. As well as measuring all of these statistics empirically, it is valuable to compute them analytically to estimate how these methods will behave in situations different than what we were able to test explicitly. The proposed methods will be evaluated using query time, percentage identity, and bit score. The percentage identity is a measure of how many characters the matched sequences share, while the bit score indicates the size of the database required to make a similar match by random chance. Generally, we are interested in the average case match that is given as well as the best and worst values returned by the queries. To this end we will be looking at the maximum and minimum values of the bit scores as well as the average returned value across queries of all query methods. Percentage identity is used as an additional indicator of the quality of the returned matches. As these values are already scaled between 0 and 100 we are more interested in the shape of the distributions. Since we must first partition the data and build the ball cover, we will also be investigating the build time for the needed indices.

## Algorithm

EnTrance explored several implementations of creating entropy scaling indexes and structures. The various clustering algorithms all choose a specific set of ball centers first and then sequences as points are added to the closest ball.

### Measuring Metric Entropy and Fractal Dimension

*Metric entropy* (21, 22) (also called Kolmogorov entropy (23)) is a measure of the diversity of a set of points in a metric space. Given a metric *d* and a radius *r*, the metric entropy of a set of points *R* is equal to the minimum number of balls of radius *r* needed to cover the set. More formally, the metric entropy *M*_r_(*R*) = min_*CϵBC*(*R*)_|*C*| where *BC*(*R*) is the set of all possible ball covers of *R* using balls of radius *r*.

*Fractal dimension* (24) (also called Hausdorff Dimension) is a measure of how the metric entropy changes with respect to the ball radius. Formally, it is the exponent *D* in the expression *p* = *r^D^* where *p* is the number of points from a data set that is covered by a ball of radius *r*. Well clustered spaces, such as biological sequences, usually exhibit a high fractal dimension.

Prior to creating the ball cover, it is helpful to be able to approximately measure the metric entropy and fractal dimension of a data set. Doing these measurements without having to construct an entire minimal ball covering can facilitate optimizing the parameters for all the other experiments. To achieve this, the Approximate Metric Entropy Measurement (AMEM) algorithm (Algorithm 2) was created to estimates the metric entropy and fractal dimension. AMEM starts with a range of candidate ball radii *R* = {*r*_1_,*r*_2_, ···, *r_n_*}. Here, ||*R*|| = 20 by default. Then, given a set of data points *P*, AMEM samples a small number *B* (set at runtime, default 100) of data points as ball centers, that is, *S*′ ⊂ *P* and |*S*′| = *B* (Algorithm 2). It then measures how many points are in the balls defined by the ball centers *S*′ and candidate ball radii. It reports the average and standard deviation of the number of points in each ball for each radius. Because this is only done with a small number of fixed ball centers, AMEM is relatively fast, but can still give a good indication of the metric entropy and fractal dimension of a data set.

**Algorithm 2.**
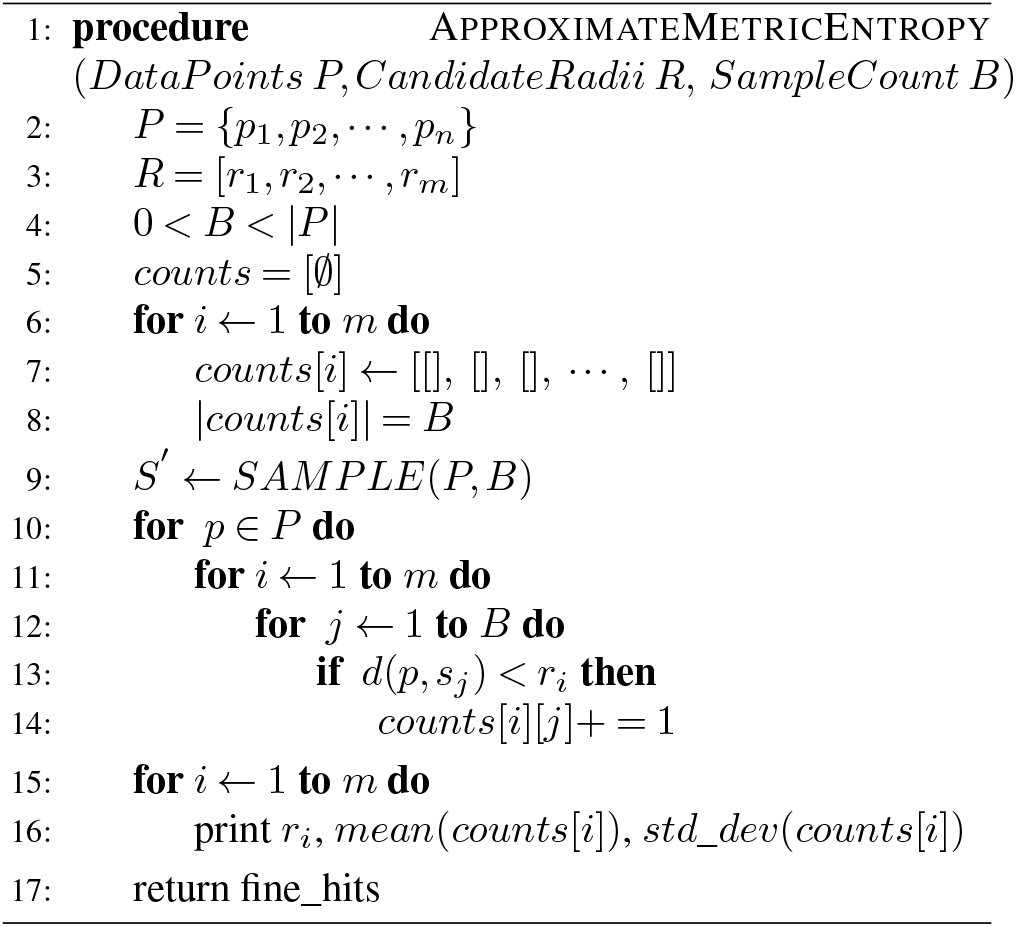
Pseudocode for the approximate metric entropy measurement algorithm. the *SAMPLE*(*P,n*) function returns a set of *n* randomly chosen elements from the set *P*.

### Random Ball Cover Construction

EnTrance randomly selects a fixed number *f* of ball centers within a given dataset. The number of balls *f* is set to the square root of *n* as default, and the overlapped ball canters are excluded. Once centers of balls are selected, randomized clustering algorithms take the dataset and add each sequence point *p* to the ball corresponding to the closet ball center *c* to minimize the distance *d*(*p, c*) where *d*(*p, c*) is under the threshold radius *r*. The point *p* is added to the temporary waiting list if there is no ball center within the radius threshold *r*. Once scanning for points in the dataset is done, points in the waiting list will be passed to this algorithm again for further iterations, by consecutively adding points into centers list. From the second iteration, a relaxed ball radius will be allowed to ensure all points will be covered after reasonable iterations. The total time complexity can be expressed as *O*(*nE*), where *n* is the number of data points and *E* is the metric entropy of the data set. In the worst case, when each point is a ball center, *E* would go to *n* and the maximum runtime is *O*(*n*^2^). But because all balls are created at the beginning, multi-workers who are employed by goroutine concurrency control in the Go language can significantly reduce the real runtime. Initially, EnTrance provides two ball cover construction: random ball cover construction and naive ball cover construction. While random ball cover construction starts construction balls with a fixed number of centers, naive ball cover constructions start with the longest sequence and add sequences consecutively into the center if the target sequence is within the threshold or updates into list of centers otherwise. During the ball cover construction on Uniref(7), we found out that naive ball cover construction results in unevenly distributed ball construction where the majority of sequences from the database are assigned to one ball. EnTrance tried to resolve this issue by randomly choosing starting sequences and changing threshold, but could not resolve the unevenly distributed construction. It remains for future work to find an optimal ways to choose the most efficient ball center sequences.

**Algorithm 3.**
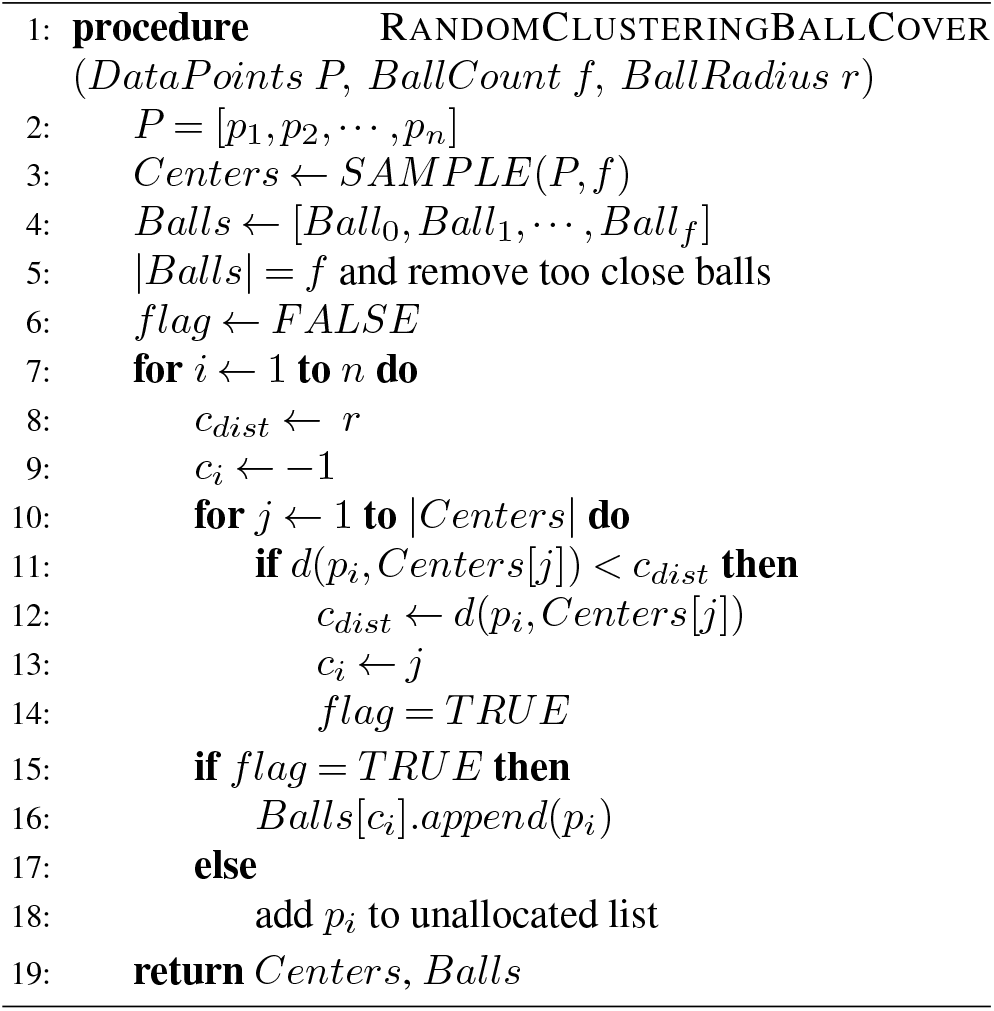
Pseudocode for the random clustering ball cover construction algorithm. The *SAMPLE*(*P,n*) function returns an array of *n* randomly chosen elements from the set *P*

### Six-Frame Translation

The prerequisite for the EnTrance entropy scaling search is translating the query DNA sequence to proteins. A given double-strand DNA may have six reading frames from forward and reverse strands and each strand has three codon loci. The translation script is modified based on GitHub repository (25). The default version of EnTrance takes protein sequences and the deployed version of EnTrance in FunGCAT(26) is set to take DNA sequences as input instead of protein sequences. This manuscript primarily focuses on sequence similarity search in protein sequences.

### Fine Search and Smith-Waterman Score

Once a coarse search generates potential balls on each length based binning, EnTrance uses BLASTP to find the sequence match. We defined this process as *fine search*. Since a fine search is done within the size of a ball, the running time for each BLASTP search is optimized. Returned sequences from a fine search will be ranked by Smith-Waterman algorithm for local alignment against the query sequence. When results sequences from fine search in each length based binning are done in parallel, EnTrance merges top matched sequences of each length based bin and writes top results in the output file.

### Parallel Computation Work

Since coarse and fine searches of EnTrance are occurring independently after length based binning, multi-workers which are employed by Go routine parallelism control in the Go language can significantly reduce the real run time. EnTrance builds and searches within length based binning independently. Once Uniref(7) dataset is distributed with length based pre-clustered ball cover construction from section 2.2, EnTrance constructs entropy scaling index for parallel sequence search. As shown in the diagram(Figure 2), once index construction is done, the input query will be searched in entire length based bins concurrently.

## Results

All testings are done on a local high-memory computation server with XeonE5620CP with 32 cores and 192GB memory.

### Test Data

All of the biological data used in these tests came from the UniProt Consortium (27). EnTrance specifically used the UniRef100 (7) protein clusters version 2018_01, which contains approximately 133 million sequences varying in length from 2 amino acids to 38105 amino acids. The current 2019_04 version contains 96 million sequences. UniRef90 protein clusters are compressed from UniRef100 by clustering the sequences in CD-HIT (28) which have sequence identity over 90%. For each of these 20 data sets, a corresponding random data set was also synthesized. Each of these synthetic data sets was created to have the exact same number of sequences with exactly the same lengths as the biological data sets, just the contents of the sequences were uniformly random over the protein alphabet. We took a stratified random sampling approach for choosing the sequences. All of the test queries consisted of single sequences randomly chosen from the largest data set and testings are done for a sample input dataset. We took *x* amount of random sequences, where *x* = 1, from every 10,000 sequences for each individual run. After choosing the sequence we took continuous partials of the sequence with a minimum partial length of 20. The minimum length is in place because small partials are not useful for testing. We take partials because the a common use case for EnTrance invovles finding novel sequences and taking partials of known sequences is a way to simulate this. EnTrance provides both sampling script to generate sample input files as we described above so that EnTrance can be tested on both test data settings in a more advanced environment in the future.

### Synthetic Data Entropy Comparison

Before constructing a ball cover on a set of biological sequences, it is useful to know what the metric entropy of the set is. The entropy measuring tool described in Section 3.1 was tested with the biological data sets as well as their corresponding randomized synthetic data sets. The result of these runs is shown in Figure 5. It is interesting how large a difference is made by which metric is used. Under Jaccard distance, the average number of points per ball in the biological data is notably higher than in the synthetic data for most radii. This implies that with a well chosen ball radius, the metric entropy of the biological data is notably lower than that of the synthetic data. This also shows the benefit of using a metric over a measure that does not satisfy the triangle inequality in all cases.

**Fig. 5.**
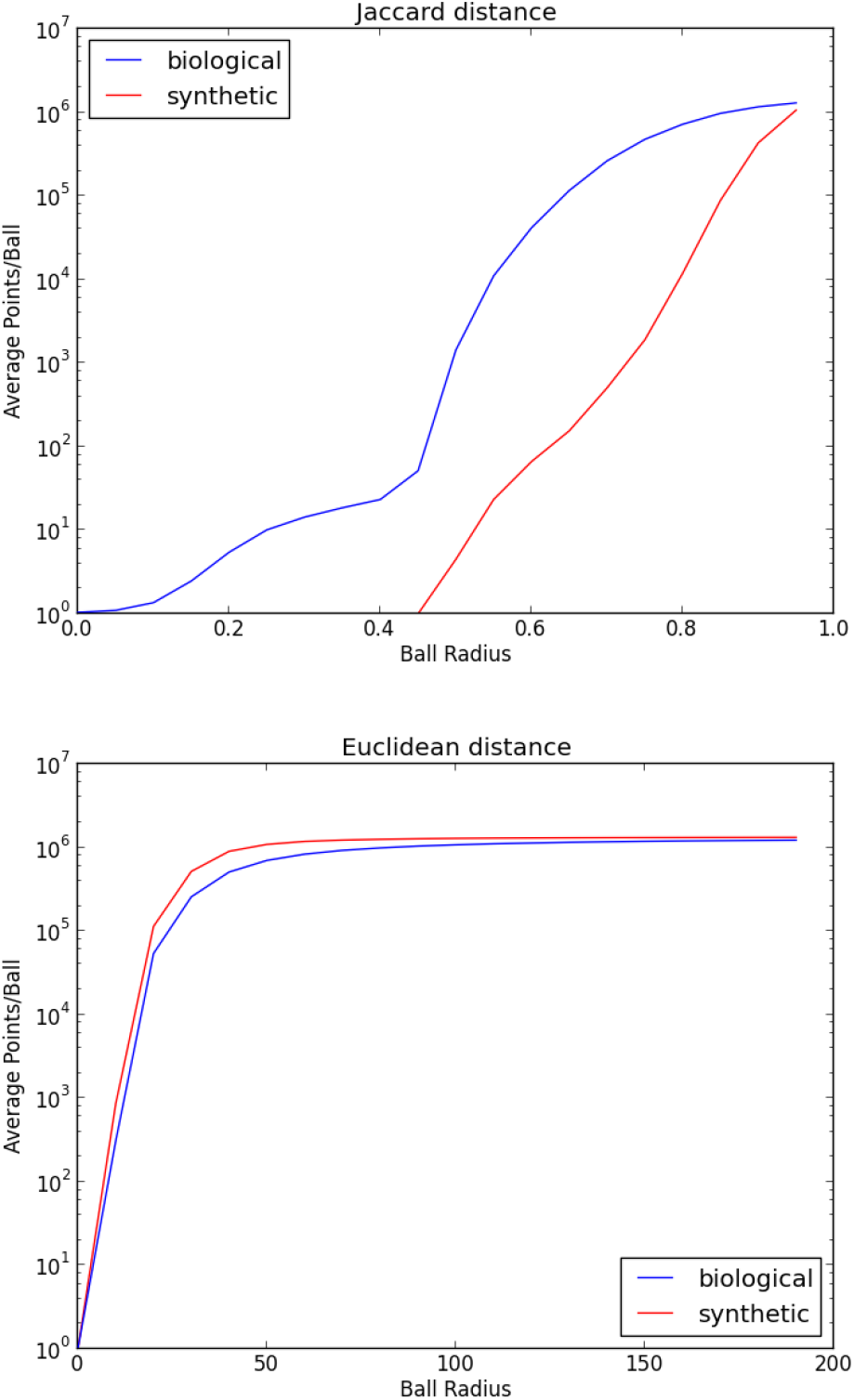
Entropy approximations of biological data versus randomized data

## Performance

### Query Time

Execution times for querying the small size sample described in the test data section are shown in Figure 6. In a given testing dataset DIAMOND performs with a total of 550 seconds as an average of 0.77 seconds per sequence. Meanwhile, a coarse search where EnTrance searches on centers database took an average of 55.14 seconds per length based bin with a range of 0.0054 to 0.016 seconds per length. Fine search ranges between a total of 2.60 to 8.27 hours per bin and 1.42 to 4.54 seconds per bin per sequence. The range could vary based on a number of input search sequences for both DIAMOND and EnTrance due to the parallelization process. Note that DIAMOND performs 20,000 times faster than BLASTX which EnTrance uses for fine search method in comparison(6). Ranges were calculated from length based bin with shortest length sequences(1 to 46) and bin with longest length sequences(1754 to 38106).

**Fig. 6.**
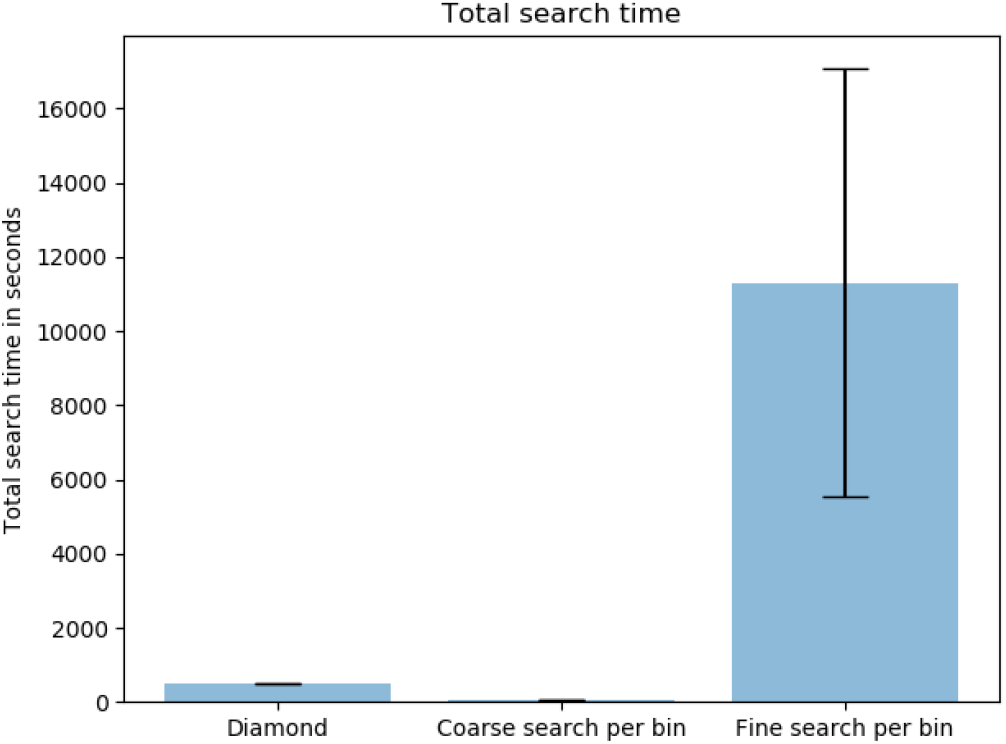
Distribution of query time for DIAMOND, coarse search, and fine search. In given testing dataset, DIAMOND performs with total of 550 seconds as average of 0.77 seconds while coarse search and fine search took average of 54.93 seconds and 13249.37 seconds per length based bin.

### Sensitivity

EnTrance principally produces more matches compared to DIAMOND since EnTrance merges fine search results from individual length based bins. Table 1 shows that EnTrance produces roughly 6.65 times more number of matches. A portion of match results of percentage identity less than 50% from EnTrance remark significantly less than those of DIAMOND. In a given testing dataset, while DIAMOND the distribution of percent identity of matches found 6860 matches with 100% percent identity EnTrance found the total number of 20174 matches. Figure 7 shows the distribution of percent identity of matches of DIAMOND and EnTrance. As we can see, matches from EnTrance consist of more sequences with higher than 60% of percent identity compared to DIAMOND.

**Fig. 7.**
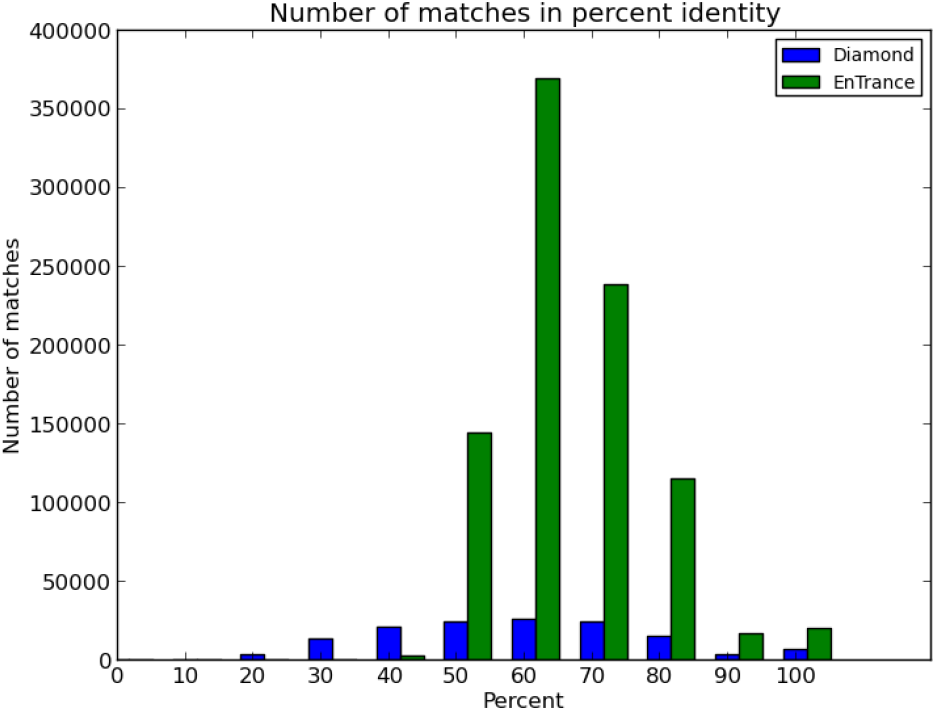
Percent identity match comparison between DIAMOND and EnTrance. Since EnTrance combines top search results of each length based bins, EnTrance overall returns more matches than DIAMOND.

**Table 1.** This table shows the distribution of percent identity in DIAMOND and EnTrance. Overall, EnTrance produces more results in general compared to DIAMOND due to merging top matches in each bin. While 45.15% of DIAMOND result has less than 60% percent identity, only 16.17% of EnTrance result is under 60%. EnTrance also produces more 100% percent identity matches than DIAMOND.

| Tool | Percent identity |  |  |  |  |  | Total |
| --- | --- | --- | --- | --- | --- | --- | --- |
|  | 0-60 | 60-70 | 70-80 | 80-90 | 90-100 | 100 |  |
| DIAMOND | 61443(45.18%) | 25907(19.05%) | 23965(17.62%) | 14891(10.95%) | 2931(2.16%) | 6860(5.04%) | 135997(100%) |
| EnTrance | 146419(16.17%) | 368756(40.73%) | 237933(26.28%) | 115108(12.72%) | 16890(1.87%) | 20174(2.23%) | 905280(100%) |

### Ball Size and Count

The entropy scaling behavior demonstrated in the query times of the EnTrance implementations is dependent on the number and size of the balls being searched. If the ball cover is well constructed, both of these quantities will be low compared to the size of the original data set. Ball indexes are constructed individually in length based bins, and EnTrance constructs the square root of the number of sequences number of balls. The number of balls per length based bin ranges from a minimum of 354 balls to a maximum of 404 balls with an average of 395.67 balls per bin.

## Discussion

In this paper, we investigated the potential usage of entropy scaling techniques in the sequence similarity search motivated by the biological sequence threat analysis problem. To do this, we started by looking at how a previous tool, MICA, tried to achieve improved query execution times based on entropy scaling. In doing this, we identified shortfalls in MICA’s design due to discontinuing in database as well as possible extensions. Based on these ideas, we explored how the measure used affects performance. The primary motivation is the tradeoff between efficiency and accuracy. On one hand we wish to maximize the number of triplets that satisfy the triangle inequality as mentioned in entropy scaling system in method section. On the other hand, we wish to use a measure that is efficient enough to scale with the considerable size of the dataset. Jaccard distance (which is not a metric) turned out to be quite efficient but at the cost of accuracy, while edit distance is a significantly inefficient metric. This directly influences the significant difficulty of index construction. Building an index that allows for entropy scaling searches can be more computationally expensive than building conventional indices and was especially complicated in situations when we were not operating with a metric. To overcome this issue, we investigated several ways to accelerate the index construction and how these methods affected query execution time and sensitivity including parallelization and length based binning. Length based binning was the most efficient choice as each bin could be treated individually, and thus in parallel at build time. This also helped address the issue that certain balls may have a disproportionate number of sequences. This did not solve the issue within each bin but was able to reduce the size of the largest ball and improve query time. Furthermore, analysis of the relative lengths of matches showed that most matches occur with sequences that fairly similar in length. This leaves an open door for further optimizations by restricting the search space to bins within a specific range. Our results indicate that while entropy scaling tools show significant promise in reducing query times compared to existing BLASTX, there is still significant work to be done in improving sensitivity and index build time. EnTrance shows more number of perfect percent identity matches than DIAMOND and detailed analysis based on length base sequences by providing logs of length based binning.

## Future Work

There are several additional ideas that have demonstrated potential, but we did not fully explore. Many other metrics and variations on the metrics we used either did not have the chance to investigate or they had a computational cost that was prohibitive to testing with large amounts of data. Using a conventional search tool, other than BLAST, that uses a metric that better aligns with whichever metric is used to construct the ball cover could significantly improve sensitivity. Entirely different ball cover construction approaches such as making use of a sequence bloom tree could significantly speed up index construction. If such an approach made it so that index construction was no longer a testing bottleneck, then the search algorithm could be tested on significantly larger data sets, hopefully demonstrating the extent to which entropy scaling would be beneficial. Another idea that would potentially improve sensitivity is to construct a network on top of the ball cover, connecting balls that are close to each other. Doing this would allow the search algorithm to search nearby balls, making it able to find less conserved sequences. It has the potential to adapt various versions of reduced protein alphabet to find optimally reduce the index building time as well. The search query can be potentially improved by selecting length based bin accordingly for input sequence length or limiting the biological domain of search query such as building a specific database of bacteria, protein, virus.

## ACKNOWLEDGEMENTS

This research was sponsored by the Office of the Director of National Intelligence (ODNI), Intelligence Advanced Research Projects Activity (IARPA), via the Army Research Office (ARO) under cooperative Agreement Number W911NF-17-2-0105, and by the National Science Foundation (NSF) grant ABI-1458359. We thank FunG-CAT team, Andrew Warren, and Phillip Anderson.

